# Gold nanorod based delivery system could bring about superior therapy effect of Ramucirumab through direct cytotoxicity to cancer cell mediated by differential regulation of phagocytosis in gastric cancer cell

**DOI:** 10.1101/2019.12.29.888537

**Authors:** Linyang Fan, Minzhi Zhao

## Abstract

Gastric Cancer (GC) is one of the most serious cancers with high incidence and mortality all over the world. Chemotherapy hadn’t led to desirable effect and targeted therapy brings about a new stage to cancer treatment. Ramucirumab is the first FDA-approved therapy for advanced gastric cancer. It is well known that gold nanorod, a nontoxic biocompatible nanomaterial, is an especially promising candidate for cancer theranostic. In this study, Ramucirumab (Ab) were first modified by gold nanoparticles to enhance uptake efficiency. The simple Nano-delivery system had taken perfect aggregation effect in vivo even better than 5-fold Ab treatment. Gold nanomaterials, especially gold nanorod (AuNR), could induce direct cytotoxic effect to cancer cell in the presence of Ab, while Ab or gold nanoparticle themselves couldn’t lead to such direct killing effect even at an extremely high concentration. Proteomic and transcriptomic analyses revealed this direct cytotoxicity derived predominantly from Ab-mediated phagocytose, and the high affinity receptor for Fc gamma CD64 showed differential up-regulation only in gastric cancer cell treated by these nanodrugs compared with Ab, especially for AuNR group. This was the first time to discover that nanoparticle could induce regulation of immune related pathways and Fcγ receptor in the target cancer cell. Simplified and powerful designs of smart nanoparticles are highly desired for clinical. The dramatic enhancement of Ab accumulation with simple composition, combined with direct cytotoxic effect specific to cancer cells brought perfect therapeutic effects in vivo than Ab, which would promote further clinical application of gold nanorod in the diagnosis and therapeutics of gastric cancer.

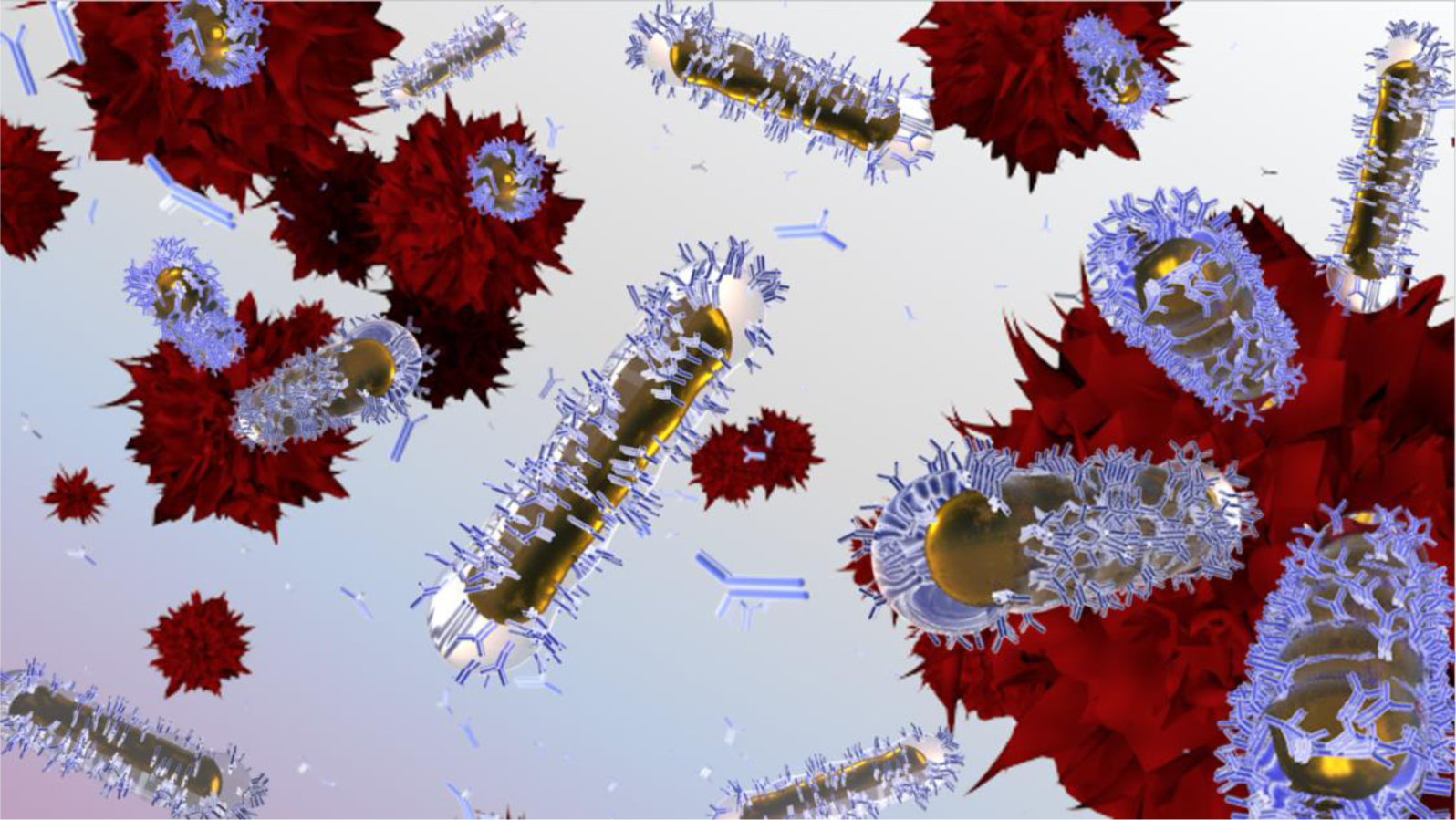

## Introduction

The Gastric Cancer (GC) is one of the most severe malignant cancer with a high morbidity and lethality in China^1^ and all over the world^2^ GC has been top 5 morbidity in USA for more than 40 years. 5-year relative survival rate was merely 30%^3^. In China, the incidence and mortality reached the 2nd among all kind of tumor. Unfortunately, most diagnosed GC patients had been already in advanced and inoperable status^4^ and chemotherapy would be the last hope for them. However, it is an undeniable fact that 5-year relative survival rate was actually double by modern diagnosis and tumor therapies development in 40 year since several gastric relative targets had been discovered, like Epidermal growth factor receptor (EGFR) ^5^, Human epidermal growth factor receptor 2(HER2)^6^, Vascular endothelial growth factor(VEGF) family ^7^, Mesenchymal-epithelial transition factor(MET)(8) and Programmed death1(PD1): PDL (PD-ligand) 1/PDL2 pathway^8^. But among these targeted drugs described above, Cetuximab (anti-EGFR)^9^, Rilotumumab (anti-cMET)^10^, as well as Keytruda (anti-PD-1)^11^ were all failure in clinical research, while ramucirumab (anti-VEGFR2) and trastuzumab(anti-HER2) were the merely approved drugs and effective to GC in the world^12,13^. And Ramucirumab had advantages in trastuzumab resistant patients^14^. Ramucirumab targeted to VEGFR2, a member of VEGF family and had a better prognosis in REGARD trial. According to this trial, patients who received ramucirumab plus chemotherapy had a median overall survival of 9.63 months compared with 7.36 months for those in the control^12^. Hence, antibody-drug conjugate (ADC) has been considered as a better therapy method^15^.

Nanomaterial-based drug delivery has been widely applied in biomedical research field including photothermal therapy, imaging and drug delivery^6,16^. It was feasible to achieve high functional ligand densities on the surface for targeting purposes because of the high surface area to volume ratio of nanoparticles^10^. Utilize functionalized nanoparticles with biomolecules that target unique or overexpressed biomarkers in tumor cells to enhance delivery efficiency. Gold nanoparticle, as the well-known nontoxic biocompatible metal, was an especially promising candidate for cancer theranostic ^17^. Due to tunable localized surface plasmon resonance (LSPR), gold nanorod are not only attractive probes for cancer cell imaging but they can also become highly localized heat sources when irradiated with a laser through the photothermal effect and can be used to provide hyperthermal cancer therapy, as well as to trigger drug release for chemotherapeutics. Thus, gold nanorod (AuNR) was selected for major investigation in the nano-delivery platform^18,19^, and gold nanosphere (AuSP) was set as control.

To establish a more efficient drug delivery platform, gold nanorod (AuNR) was chosen, linking by polyethylene glycol (peg) to antibody Ramucirumab (Ab) and a chemotherapy drug Doxorubicin (Dox), which was a chemotherapy drug and applied in GC for more than 20 years^20,21^. The preparation process of AuNR-peg-Ab-Dox was showed in **Scheme 1**. The physical and biological effects of these nano-platforms were evaluated *in vitro* and *in vivo* respectively.

**Scheme 1.**
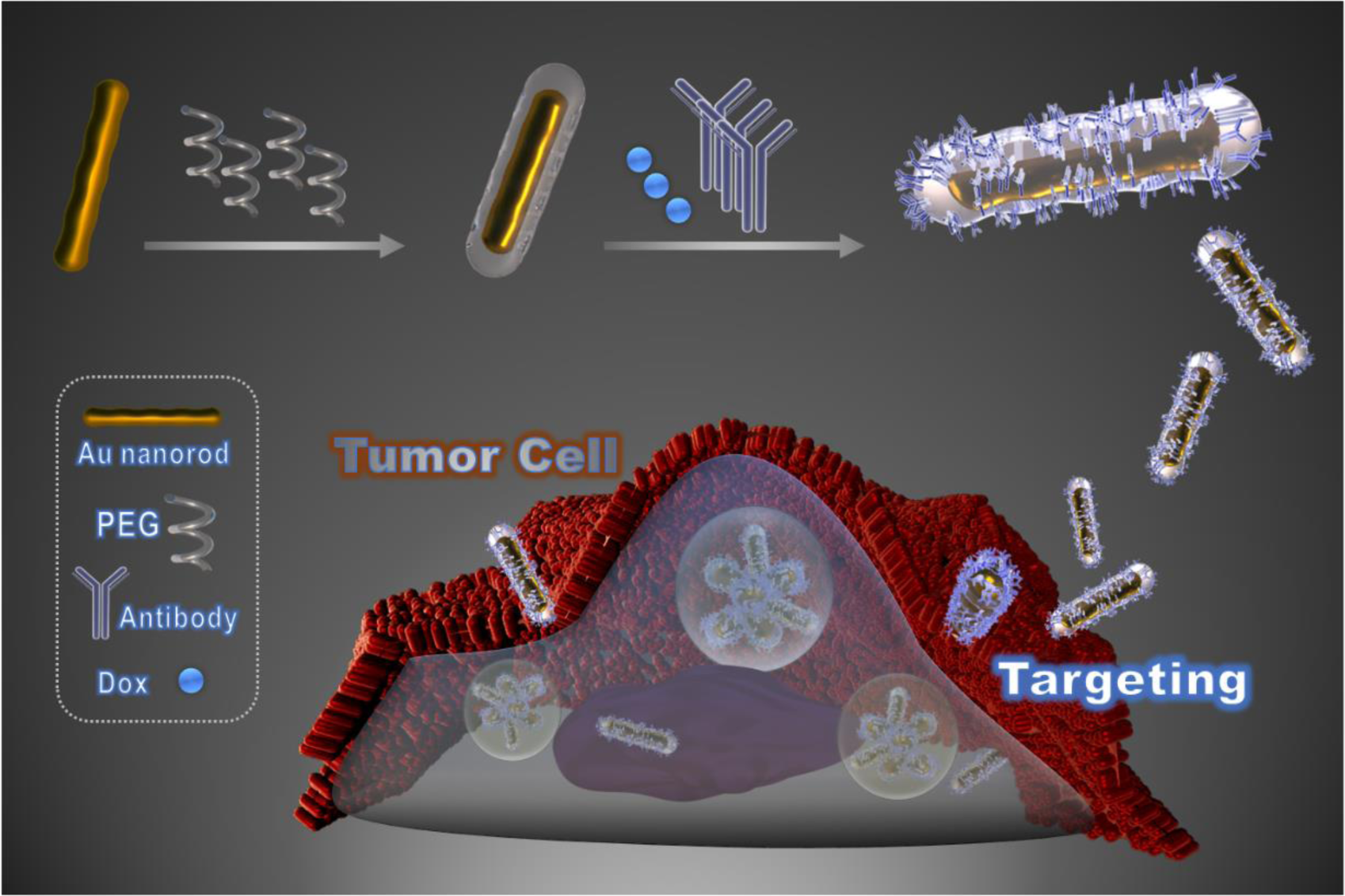
The fabrication and drug delivery of Au-peg-Ab-Dox.

## Results and Discussion

### Synthesis and characterization of gold nanomaterial delivered drugs

The composite materials were characterized (**Figure 1**). There were obvious capsules on the surface of gold nanorods (AuNR) and nanosphere (AuSP) characterization by TEM (200000×), which confirmed the successful coating on the surface of the gold nano-particles (**Figure 1 A-B**). The capsule of nanoparticles was thicker when antibody was added into materials. **Figure 1 C-D** showed that the synthesis of all kind of materials was accomplished by spectrophotometer detection. And spectrum exhibited red shift when Au modified by Peg. When Au-peg linked to Ab, the sharp of spectrum changed.

**Figure 1.**
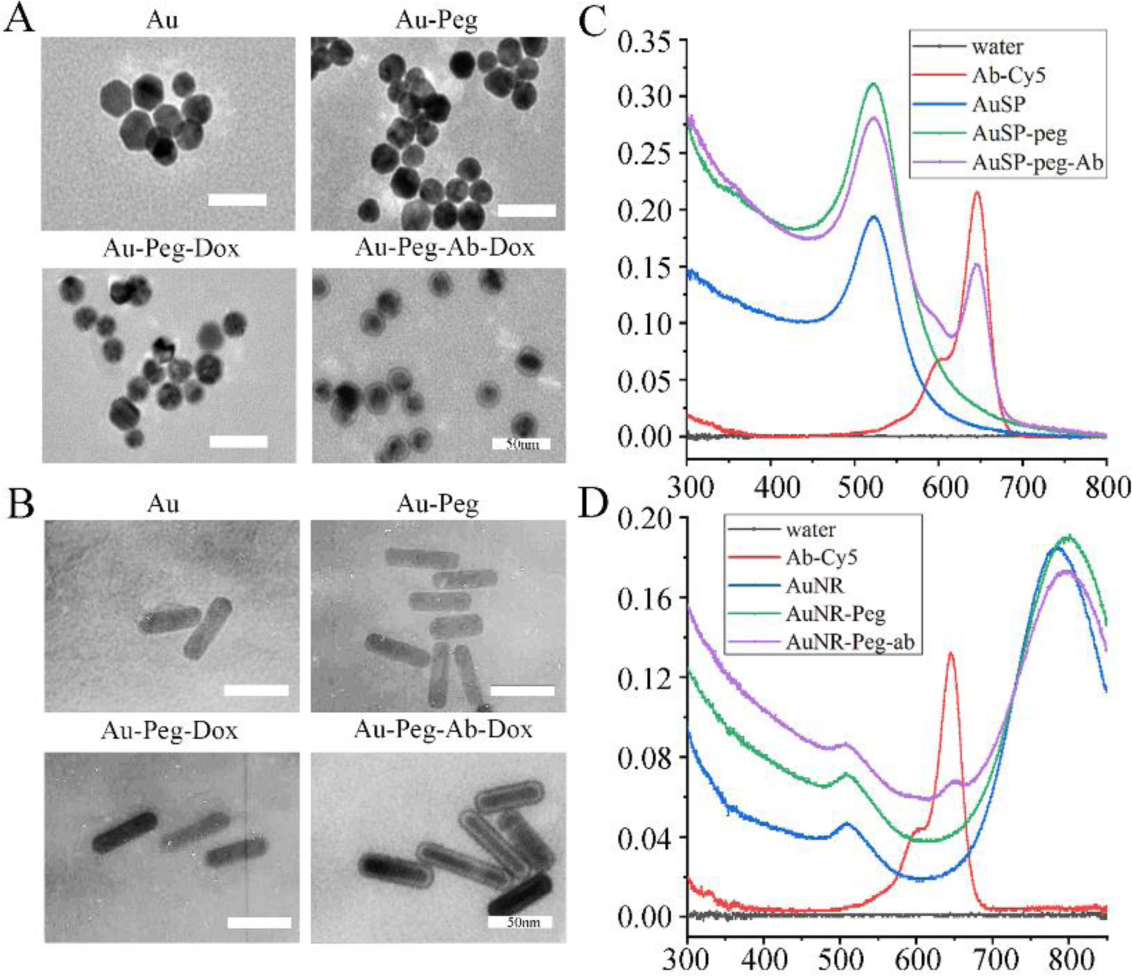
Gold nanoparticles characterization by TEM and spectrum. A, C) TEM detection for gold nanosphere and nanorods (AuSP and AuNR), Au-peg, Au-peg-Dox and Au-peg linked by Dox and Ramucirumab (Ab). B, D) Spectrum measured red shift when Au modified by Peg and Ab.

### In vitro and in vivo imaging for the uptake and accumulation efficiency of nano-drugs

To evaluate the tumor cells recognition, combination speed and the uptake efficiency of nanoparticles, a low concentration of antibody (1 μg/mL in Ab, FDA recommend 8mg/Kg, equal to 8μg/mL) was applied, and confocal laser scanning microscopy (CLSM; 63×, oil-immersion objective) imaging and flow cytometry (FCM) were employed (**Figure 2 A-E**). GC cell SNU-5 were treated by Ab, peg-Ab, AuSP-peg-Ab, or AuNR-peg-Ab (1 μg/mL Ab) for 1, 2, 4 hours. The fluorescence in cells was increasing from Ab to AuNR-peg-Ab group and cells treated by Ab for 4 hours were still appeared vague. The similar phenomenon was exhibited in AuSP-peg-Ab and AuNR-peg-Ab treated cells, as the triple fluorescence intensity than Ab by FCM measurement at 1hour. In fluorescence imaging results, 2 hour’s treatment of AuNR and AuSP performed equally to peg-Ab treated for 4 hours and much better than Ab. What’s more, AuNR and AuSP contained nanodrugs were obviously endocytosed by cells in 4 hours, as Ab and Peg-Ab were still stained encircle the cells. To test the gathering effect of Ab *in vivo*, all drugs were intravenous injected into tumorigenic BALB/c-nu mice (**Figure 2F-H**). AuNR-peg-Ab was injected at 30μg Ab, meanwhile Ab at 30 μg and 5× at 150 μg according to human recommended dose 8mg/kg ^22,23^. No tumor mouse injected by Ab showed that the majority of drugs were metabolic located in liver. To evaluate drug recognition ability, fluorescence was measured in tumor and normal tissues by Image J software. The biodistribution of the excised organ fluorescent images showed that Ab (30 μg) performed poor and there was no significant change of fluorescent intensity until the quality of Ab increasing to 5-fold. AuNR-peg-Ab (30 μg) performed better than Ab (30 μg) and 5×Ab (150 μg) group. AuNR-peg-Ab group provided similar performance in all groups at 2 hours detection, while there was a significant increase at 8 hours and max fluorescent intensity at 24 hours in AuNR-peg-Ab group (**Figure 2G**). What’s more, AuNR-peg-Ab’s F_ratio_ raised 10 times than non-tumor and Ab (30 μg) in 24 hours (**Figure 2H**).

**Figure 2.**
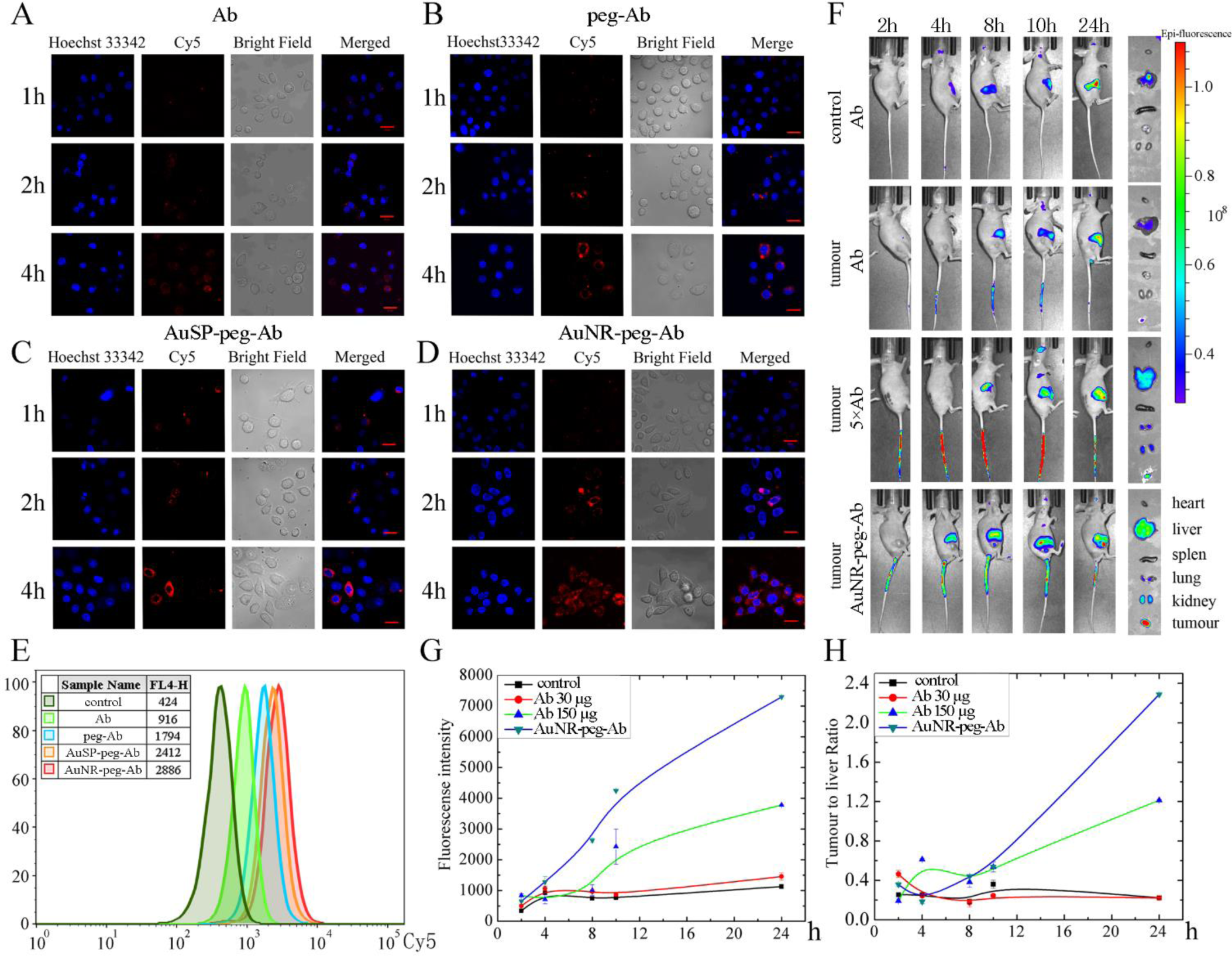
A-E) Combination detection of Ramucirumab (Ab) linked gold nanoparticles in SNU5 cells. A) Ab treated SNU5 cells. B) Ab linked to peg treated SNU5 cells. C, D) Gold nanoparticles (AuSP and AuNR) linked Ab treated SNU5 cells. E) Flow cytometry analysis of nanoparticles cellular uptake efficiency. F-H) In vivo imaging of the gold nanomaterial in the SNU5 tumor bearing mice. F) Images of tissue and major organs at different time periods after intravenous injection of Ab and AuNR-peg-Ab. G, H) fluorescent intensity (F_tumor_) and F_ratio_ (F_tumor_ to F_Liver_ ratio) measurement at different time.

### Analysis of toxic effects of different nano-carriers and nano-drugs at cellular level

VEGF/VEGFR2 (KDR) is a classical interaction signal pathway in HUVECs^24^. VEGF induced HUVECs to overexpress VEGFR2 and promote angiogenic sprouting, endothelial cell differentiation and permeability directly^25-27^. Besides, the VEGF/VEGFR-2 system played an important role in survival, proliferation and anti-apoptosis in many kinds of cancer cells^28^. We used 20 μg/mL VEGF to stimulate HUVECs as a VEGFR2 reliable cell model and VEGFR2 cell affinity validation assay showed cell toxicity equaled to 6.6 μg/mL(C value in 4 parameter curve) and fulfilled FDA requirement(8 mg/kg), indicating that Ab could meet the need of quality control (**Figure 3A**). However, Ab cannot provide equal effect for SNU5, MKN-45 (another gastric cancer cell) and GES-1(normal stomach cell) in our research (**Figure 3B-D**). In detail, SNU5, MKN-45 and GES-1 cellular activity had no change in 10^−2^-10^3^μg/mL Ab, whereas HUEVC cell results formed a normal S sharp. In a word, the Ab treatment alone hadn’t taken significant influence to tumor cell viability.

**Figure 3.**
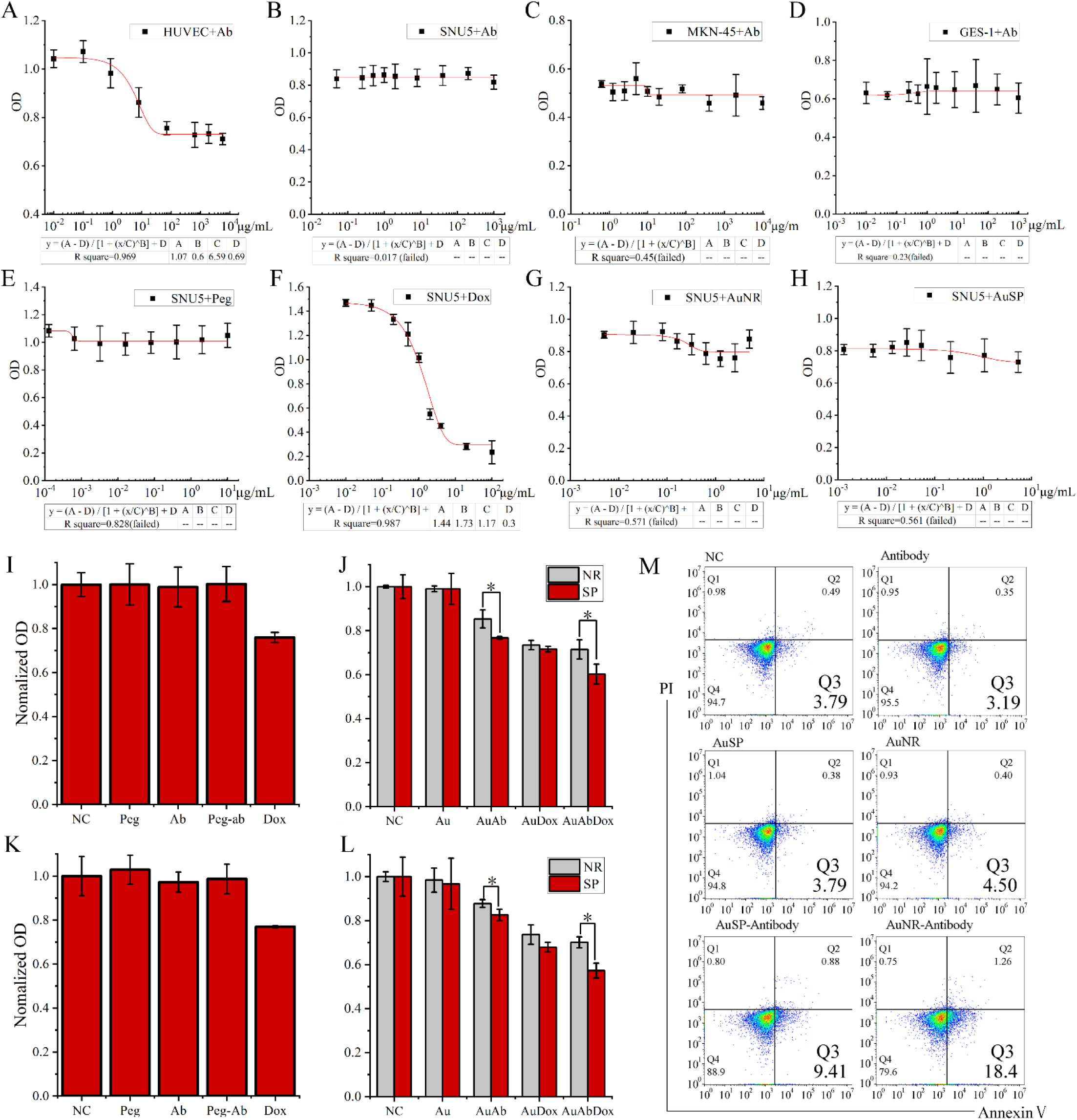
A-D) IC_50_ curve treatment by Ramucirumab (Ab) in HUVEC cells (A), SNU5 cells (B), MKN-45 cells (C), and GES-1cells (D). E-H) IC_50_ curve treatment by Peg (E), doxorubicin (Dox) (F), AuNR (G) and AuSP (h) in SNU5 cells. I-J) MTT measurements of cell viability in SNU5 cells treated with different components. *\*, p*<*0.05*, n=6. G-H) MTT measurements of cell viability in MKN-45 cells treated with different components. *\*, p*<*0.05*, n=6. M), cell apoptosis analysis by flow cytometry. Peg was omitted in the mark of the nanodrug complex.

Our nanoparticles contained AuNR/AuSP, Peg, antibody and Dox. AuNR/AuSP was the core of the complex. Peg played a role of linker between Au and Ab/Dox. Ab played a role of scout for searching target cells and Dox played a role of soldier for enhancing toxicity. To be specific, cell viability tests were carried out to the series of nano-vehicles for evaluating the cytotoxicity to GC cells. The results indicated that neither gold nanomaterials nor other linkers had cell inhibition effect, except for Dox (**Figure 3E, I, K**). When these components were formed as Au-peg-Dox, it inhibited cell viability prominently. It was interesting that the nano-carriers can strengthen the inhibition of cell viability when linked to Ab, no matter the existing of Dox or not. Furthermore, the promotion effect was significantly stronger in the AuNR-peg-Ab treated cells than AuSP-peg-Ab group (**Figure 3J, L, *p*<0.05**). This distinction hadn’t seen in single gold or Au-Dox group. In order to demonstrate whether this difference was only showed in SNU5 cell, MKN45 cell (another GC cell) was tested and the results were the same. In apoptosis analysis, both AuSP-peg-Ab and AuNR-peg-Ab could induce SNU-5 cells annexin V(apoptosis maker) exstrophy. Furthermore, Q2 cell ratio by AuNR-Ab was more than AuSP-Ab, hinting that the different shape of material caused diverse physiological effects (**Figure 3M**). It was the first time that we found AuNR based nano-carriers had shape-dependent inhibition of gastric cancer cell viability in the presence of Ab. The recent report for toxicity was not for sphere or rod shape^29^. The earlier studies on shape-dependent toxicity of gold nanomaterials which were focused on spherical- and rod-shaped nanoparticles did not discussed the shape ^30-33^. The only one for shape effect of toxicity which indicated that spherical gold nanoparticles were generally more toxic than rod-shaped particles suggested that the higher toxicity of CTAB-coated spheres came from a higher release of toxic CTAB upon intracellular aggregation^34^. Our results were different from all investigations discussed above. The coating used in this study was Peg, which was regarded as biocompatible and its toxicity was also confirmed in our results (**Figure 3E, I, K**). And Peg-Ab also showed nearly no toxicity (**Figure 3I, K**). In addition, the AuNR and AuSP themselves, as well as combined with Dox, showed no difference on toxicity in two GC cells. Thus, this shape-depend cell toxicity had close relation with Ab.

Our results had proved that the intracellular concentration of Ab in GC cell was nearly the same for both AuNR-peg-Ab (NR group) and AuSP-peg-Ab (SP group) (**Figure 2C-E)**, as well as Ab itself didn’t influence the cell viability/toxicity (**Figure 3A-D**). Although the existing of gold nanomaterials enhanced the accumulation and uptake of Ab, but the most aggregation enhancement came from Peg (**Figure 2B, E**), which hadn’t induced inhibition of cell viability as Peg-Ab (**Figure3 I, K**). Then it could be deduced that the reason for the difference might not mainly come physically from the mount of Ab, but from the biological interaction of gold nanomaterials with Ab.

### Quantitative proteomics and transcriptomics analysis for the mechanism of the cytotoxicity diversity between NR group (AuNR-peg-Ab) and SP group (AuSP-peg-Ab)

In order to discover the biological mechanism for the difference of GC cell viability at molecular level, mass spectrometry based quantitative proteomics and RNA-sequencing based transcriptomics strategies were employed on NR group (AuNR-peg-Ab) were compared with SP group (AuSP-peg-Ab). The results were shown in **Figure 4.**

**Figure 4.**
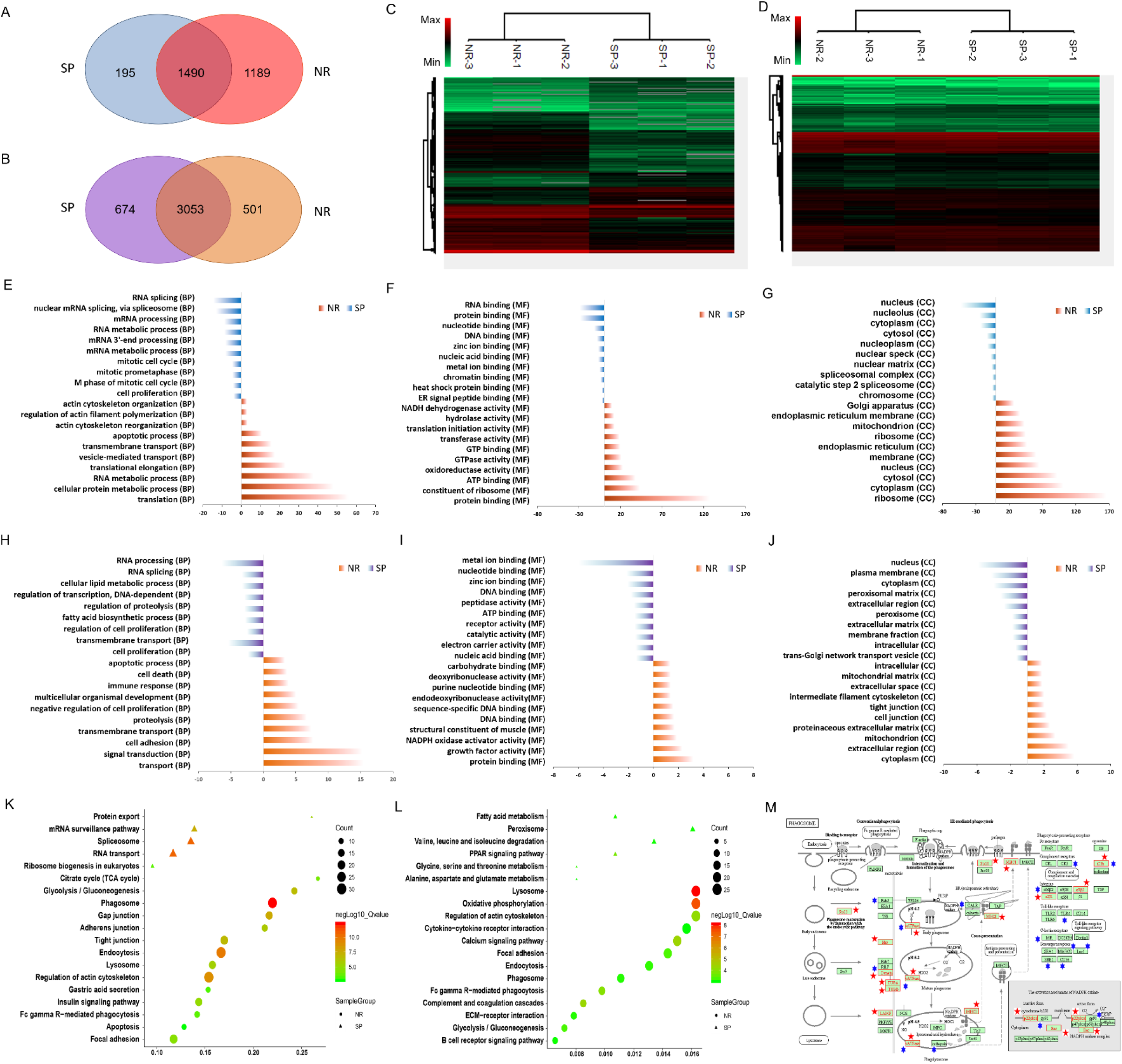
Quantitative proteomics and transcriptomics analysis of AuNR-peg-Ab (NR group) compared with AuSP-peg-Ab (SP group) in SNU5 cells. A, B) Venn grams of the whole numbers of differential proteins and expressed genes quantified in proteomics and transcriptomics from different groups. C, D) Differential protein and expressed gene heat maps for NR and SP groups. E-G) Top 10 items in the Gene Ontology (GO) biological process (BP), molecular function (MF), and cellular component (CC) enrichment of differential proteins in the proteomic results. H-J) Top 10 items in the GO BP, MF, CC enrichment of differential expressed genes in the transcriptomic results. K-L) KEGG enrichment of differential proteins and expressed genes. the color indicated the level of q value and the items with q<0.001 were included. M) Pathway “phagosomes” from KEGG with altered proteins (red pentagram) and expressed genes (blue hexagram) mapped in it.

The number of altered proteins and expressed genes only in NR group were 1189 and 501, while 195 and 674 in SP group, with 1490 proteins and 3053 genes overlap (**Figure 4A-B**). Although with similar number of altered genes, the number of altered proteins was obviously more in NR group. Hierarchical clustering for these differential expression protein and gene sets was created with an absolute log2 fold-change cutoff of ≥1 and an adjusted p value (q value) of ≤0.05. Large amount of proteins and differential expressed genes were derived from the different nano-drugs. It can be seen that the biological replications were clustered together and different groups were separated well. In addition, the pattern of the mRNA level was relative uniformity for NR and SP group, while in the protein level the difference was more apparently. Furthermore, the functional enrichment was analyzed via Gene Ontology (GO) biological process (BP), molecular function (MF), cellular component (CC) and Kyoto Encyclopedia of Genes and Genomes (KEGG) pathway. The proteins and genes which presented only in and up-regulated in NR group were recognized as NR altered, and the ones which presented only in and up-regulated in SP group were recognized as SP altered. **Figure 4E-G** showed the top 10 enriched items in GO o proteomic results and **Figure 4H-J** showed the top 10 enriched items in GO from transcriptomic results. There were several items displayed similar in both protein and mRNA level. For example, apoptotic process (BP), transport (BP), and protein binding (MF) appeared in NR group. RNA splicing (BP), cell proliferation (BP) metal ion binding (MF) and nucleus (CC) appeared in SP group. Together with cell death (BP) in NR group and cell cycle related process of BP in SP group, it showed that NR group possessed more inhibition effect to cell viability relative to SP group. It can be seen that translation (BP) showed predominantly enrichment in proteomic results, which suggested that there were many differences coming from translation, the step after transcription. This can also explain the reason that there were comparable differential expression genes for both NR and SP group in transcriptomic results, but more proteins distinct for NR group. The apoptotic process (BP) enriched in NR group, which displays one source of the more inhibitory effect of AuNR to tumor cells.

Then the enrichment for KEGG signal pathways were analysis from proteomic and transcriptomic level (**Figure 4K-L)**. There were more pathways enriched in NR group than SP group for both levels. “Phagosome” and “Lysosome” were the most enriched categories at proteomic and transcriptomic level. Phagocytosis plays an essential role in host-defense mechanisms through the uptake and destruction of pathogens ^35^. Cross-linking of Fc of Ab initiates a variety of signals mediated by tyrosine phosphorylation of multiple proteins membrane remodeling to the formation of phagosomes. Nascent phagosomes undergo a process of maturation that involves fusion with lysosomes.^36,37^. “Phagosome” showed obvious enrichment at both proteomic and transcriptomic level, and the members enriched in this pathway were labelled in **Figure 4M.** Some other immune related items like “Fc gamma R(FcγR)-mediated phagocytosis” which related with antibody directly, as well as “Complement and coagulation cascades” and “B cell receptor signaling pathway” were also enriched in NR group. Multiple mechanisms have been observed for cancer therapy by monoclonal antibody, including the Fc-dependent effector mechanisms, complement dependent cytotoxicity (CDC), and Ab-dependent cellular cytotoxicity (ADCC) ^38^. The FcγR, B cell and complement related pathways were enriched, which indicated the relation of difference had certain correlation with Ab function and immune response, which also showed in GO BP result of NR group at transcript level.

### Gold nanocarrier induced the direct cancer cell cytotoxicity by the immune related phagocytose mediated by differential regulation of FcγR CD64 on cancer cell

However, there were only tumor cells existed in this study. It was reported that antibody may induce programmed cell death (PCD) directly correlated with reactive oxygen species (ROS). This pathway evokes PCD directly in the target cell in an actin-dependent, lysosome-mediated process. ROS generation was mediated by nicotinamide adenine dinucleotide phosphate (NADPH) oxidase^39^. Antibody mediated PCD pathway can be enhanced via Fc cross-linking by secondary Antibody or FcγR-expressed cells^38^. These features were fitted to the significant enriched pathways in our study, like “Phagosome”, “Regulation of actin cytoskeleton”, “Lysosome”, “Oxidative phosphorylation” and “Endocytosis”. Some categories in the GO BP and GO MF enrichments also had relation with this, like “NADPH oxidase activator activity” ^40^. ROS generation was also detected (**Figure 5A**). The results showed that the ROS value in Au-peg-Ab was higher than individual Ab or Nano-materials. Furthermore, NR group presented higher value than SP group, which could be one of the reasons for a higher cell death rate of NR group.

**Figure 5.**
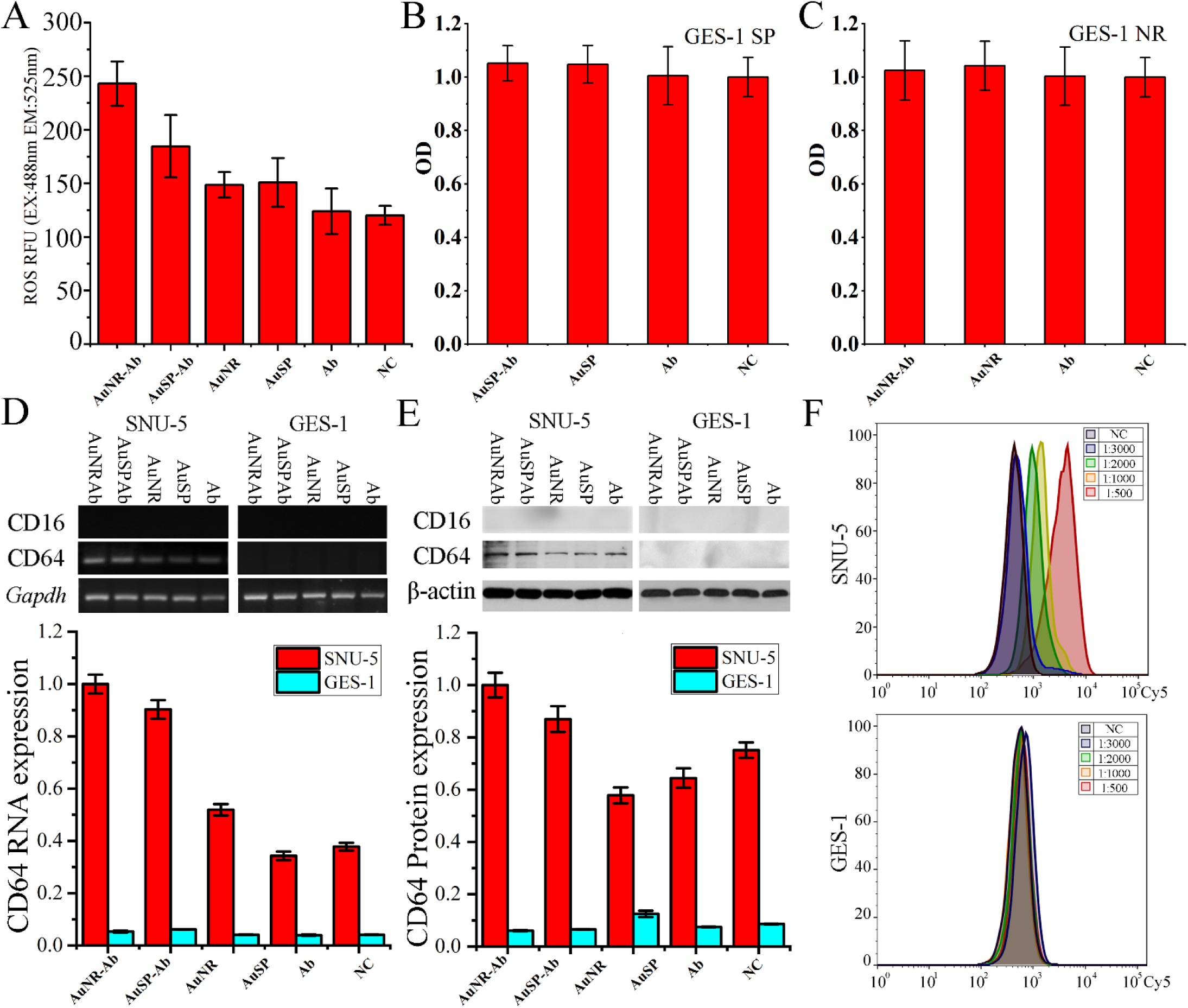
A) ROS generation test of SNU5 cell. B-C) Cell viability analysis in GES-1 cell treated by different material. D-E) mRNA and protein expression level CD64 in SNU-5 and GES-1 cells. F) Flow cytometry analysis of Ab in SNU-5 and GES-1 cells. Peg was omitted in the mark of the nanodrug complex.

Except for professional phagocytes including macrophages, neutrophils, and monocytes, epithelial cells, fibroblasts and other cells can take up particles ^41^. Phagosomes can also be produced in non-professional phagocytes. Since fibroblasts lack Fc receptors, the antibodies could not act as an opsonins but only take effect through a common ‘non-immunological’ mechanism. The cell type used in the mechanism study was tumor cell, not fibroblasts or epithelial cells. There isn’t any report said that tumor cell was phagocyte. Even if tumor cell could also be regarded as non-professional phagocytic cell, tumor cell hasn’t been realized to possess immune related function or has immunoreceptor. However, it could be seen obviously about the immune process and immunoreceptor related enrichments in our proteomic and transcriptomics results, like “Phagosome” and “Fc gamma R(FcγR)-mediated phagocytosis”. Thus, the expression levels of Fc gamma receptors, CD64 and CD16 ^42^, were detected from the mRNA and protein level (**Figure 5D-E left**). The results showed that CD64, high affinity receptor, had noticeable expression in tumor cells, and CD16 didn’t show any expression. The Au-peg-Ab groups which containing both Ab and nanomaterials showed significant up-regulation than the left groups. It was up-regulated in NR group compared with SP group (*p*<0.05). And the Ab treated group showed similar expression level with the groups only treated with nanomaterials. This was in accordance with the cell viability results of tumor cells (**Figure 3I-L**). Gastric cancer cell endogenously express Fc gamma receptor (FcγR) may facilitate the cytotoxicity, which might be the important reason that gave Au-peg-Ab groups higher cell death effect of tumor cell even without the help of immune system. The expression situation of CD64 in normal gastric epithelial cells (GES-1) were also been tested. The results displayed that none of these group showed any expression (**Figure 5D-E right**). We supposed that the strong endogenous expression of cancer cell was an important reason for the cytotoxicity of these Ab containing nanodrugs. Thus, we further investigated the cell viability of GES-1 cell. The results in **Figure 5B-C** verified that all these treatments didn’t possessed any cell cytotoxic effect to GES-1cell compared with control. It was interesting that the gold nanomaterials could activated the endogenous expression of FcγR specifically in gastric cancer cell when it was linked with Ab, meanwhile Ab couldn’t achieve this alone. Since SP group also up-regulated CD64 in SNU5 cell compared with Ab treated group, so the transcriptome of AuSP-Ab and AuNR-Ab were analyzed respectively, compared with Ab treated group. The results showed in Figure S4 indicated that both of them aroused up-regulations correlated with the immune related processes and the antibody directly related pathway like “Fc gamma R(FcγR)-mediated phagocytosis” and “Phagosome”. However, NR group triggered more enrichments than SP group, and NADPH related process only showed up-regulation in NR group. This is the reason that NR group showed up-regulation in these pathways and processes in the direct comparation to SP group. From the results of flow cytometry analysis to the different concentration of Ab treated SNU5 and GES-1 cells, it could be seen that GES-1 cell hadn’t internalized any Ab even at a high added concentration (**Figure 5F)**. This meant that a common ‘non-immunological’ endocytosis mechanism might not be the predominant reason of Ab. For the GC cell SNU5, it indicated that the internalizing of Ab increased with the rising of the added Ab amount. Meanwhile, the cell death effect was not influenced by it (**Figure 3B**). This meant that AuNR not only increased the uptake and accumulation of Ab, but also could induce cell death specific to cancer cell via the activation of immune receptor and immune phagocytose related pathway without the assistance of immunologic cell. The most important was that Ab must be contained but couldn’t achieve this itself even at an extremely high concentration. These results suggested the superiority of AuNR and it would be benefit for the clinical application of gold nanorod in the immunologic therapy, especially for gastric cancer (as this Ab can’t be recognized and uptake by another tumor cell like Hela, see **Figure S5**).

### *In vivo* analysis of nano-drugs in inhibiting tumor growth

On the base of in vitro tests, the immunologic therapy effect in vivo was carried out on SNU5 xenografted mouse model. 16 mice were randomly divided into four groups (n=5). They were intravenously treated with Ab (8 mg/kg), AuSP-peg-Ab (8 mg/kg in Ab), AuNR-peg-Ab (8 mg/kg in Ab) and PBS (100 μL/dose), respectively. The treatment progress lasts 14 days. The tumor volume and body weight of each mouse were measured every other day. As shown in **Figure 6A**, the tumor growth rate of the NR and SP groups were more inhibited obviously compared with the other two groups. And the Ab treated group only showed slightly higher inhibitory effect than PBS treated group. NR group displayed the best therapy effect in all of the four groups with comparable body weight (**Figure 6A**). Tumor and main organs were excised after the treatment. The hematoxylin-eosin (H&E) stained tumor from the NR group showed significant damages in tumor tissue. Besides, all the immunologic therapy groups hadn’t showed obvious damage to normal organs for the morphologies of heart, liver, spleen, lung, and kidney had no obvious difference to that of the PBS treated group (**Figure 6C**). The results were further confirmed by the terminal deoxynucleotidyl transferased dUTP Nick-End labeling (TUNEL) assay (**Figure 6D**). NR and SP groups showed obvious cell apoptosis degree, and NR possessed more.

**Figure 6.**
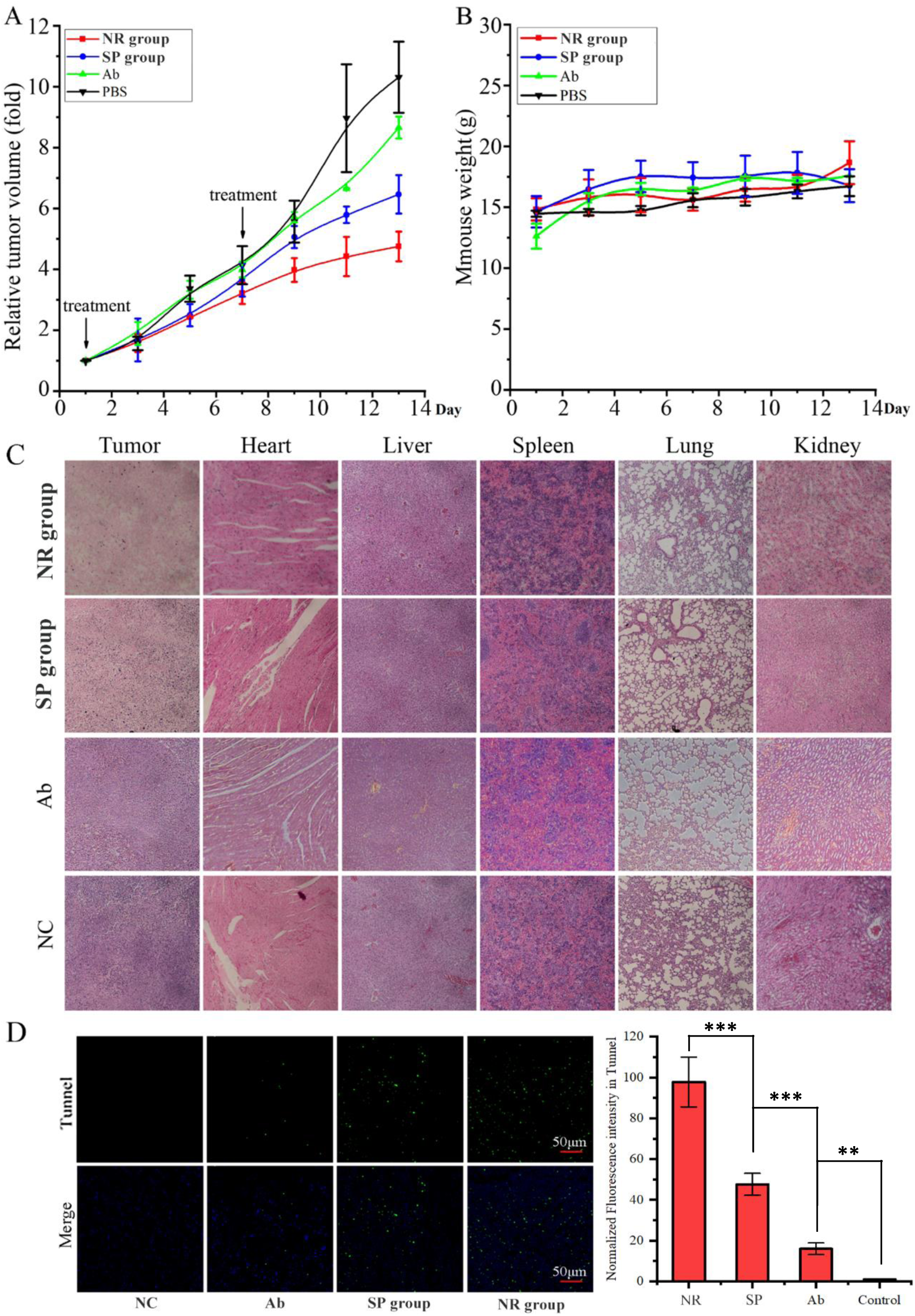
In vivo evaluation Ab connected nanomaterials (Ab, NR group and SP group) in tumor inhibition. A), The changes in tumor size during the curing process. B), The changes in mice body weight during the curing process. C), H&E images of the tumor and other tissue slices stained with H&E. D), Tunnel immunofluorescence straining detected apoptosis cells. **, p<0.01, ***, p<0.001, n=5.

These founding in this study had two meanings. First, the gold nanorod can induced stronger biological response not because carrying more antibody, but via evoking response of target cell and inducing molecular function directly. What’s more, this promotion effect can take effect only with the existing and assistance of antibody. Thus, this unique function of gold nanorod is specific to antibody-targeted therapy, which means that neither chemotherapeutic drug nor small molecular targeted drugs can get this effect. No matter for the advance of nano-delivery system, or for the advance of antibody-targeted drug themselves, these results would give more incites for further exploring in the biological effect of nanomaterial and antibody interactions.

## Conclusion

In summary, we focused on promoting the effect of the promising targeted monoclonal antibody drug Ramucirumab (Ab) for GC, by the gold nanorod with perfect physicochemical and biocompatible properties. It was found that antibody could enhance the recognition, uptake and accumulation efficiency by nano delivery system *in vitro* and *in vivo*. Furthermore, the nanodrug could induce cell death effect in GC cells directly but not in normal gastric cell, compared with Ab and nanomaterial themselves. According to proteomic and transcriptomic results, the cytotoxic effects mainly came from Fc gamma receptor-mediated phagocytosis. Furthermore, it was found that *Fcgr1*(CD64), the high affinity receptor gene for Fc gamma, was up-regulated in nanodrugs, especially for AuNR-peg-Ab group. Its high uptake and gathering, as well as direct cytotoxicity specific to GC cells explained its excellent therapeutic effect in vivo. The simple delivery system was more beneficial for the application of nanomaterial in clinic for better safety and controllability, and these findings will give more references for the better application of gold nanorod in the drug-delivery therapy in gastric cancer.

## Methods

### Design of gold particles-peg conjugate to Cy5-labeling of Ramucirumab and/or doxorubicin

Our original goal was enhance cytotoxin effeciency of Ramucirumab. Nanoparticles as an agency combined with chemtherapy and phototherapy, would synergistically abrogate tumors and prevent their recurrent, either with or without tumor resection^18^. Hetero-functional Peg (α-Mercapto-ω-carboxy Peg solution, HS-C2H4-CONH-PEG-O-C3H6-COOH, MW. 3.5 kDa) link to 9-15 nm dia, x 46-56 gold nanonanorods(Alfa Aesar46819) in 18 MEG DI water mixed with SDS for 12 h in RT, yet Peg do not affect activity in HUVEC cells. Peg excess was removed by centrifugation when Peg was quantified by the Ellman’s Assay to calculate the concentration whether Peg was all/half link to gold nanoparticles. 10kDa dialysis to remove Peg as to preparation amount of Au-Peg. Optimized ratio with gold nanorods to PEG is 1mL:0.14 mg as all of Peg link to gold nanonanorods. Lower ratio of Ramucirumab to Alexa Fluor® 647(Ex: 650 nm, Em: 670 nm) to ensure there were Cy5-free primary amines (R-NH2) of antibody. Ramucirumab and/or doxorubicin functionalized with Au-Peg by standard EDC/NHS coupling reaction^43^.220-850nm Spectrum and TEM validate that Au-Peg, Au-peg-Ab(two peg ends link to gold nanoparticles and Ramucirumab), Au-Peg-Dox(two Peg ends link to gold nanoparticles and doxorubicin), Au-peg-Ab-Dox (one end in Peg link to gold nanoparticles and the other end link to Dox and Ramucirumab) were synthesis correctly.20nmGold nanospheres(STREM 95-1547) replaced nanorods to figure out whether materilal sharp affect cytotoxin performance.

### Cell culture and cell viability analysis

HUVEC cell was purchased from Allcells (Endothelial Cell Medium, H-004B, Allcells, supplemented with 5%fetal bovine serum), GC cells SNU-5 (1640 Cell Medium, Gibco, supplemented with 10% fetal bovine serum) cell was purchased from ATCC and MKN-45(1640 Cell Medium, Gibco, supplemented with 20% fetal bovine serum) cell was obtained from ScienCell.SNU-5 and MKN-45 cells arepoorly differentiated gastric carcinoma cells.

Firstly, we established a VEGF-reliable cellular growth model in HUEVC cell(Fig S1), made VEGF protein as a stimulate factor to HUVEC cell activity. 9000 cells seed in ever well in 96-well plate with Endothelial Cell Medium. VEGF added into medium as a final concentration of 12.5 ng/mL in the next day. After 96 hours of incubation, cell was starvation by basal Endothelial Cell Medium overnight, Ramucirumab dilutedby Endothelial Cell Medium supplemented with 5% fetal bovine serum to final concentraion on a range of 10^−2^-10^4^μg/mL and mix into per well for 24 hours. The concurrent method was applied in SNU5. While doxorubicin treatment to SNU5, tumor cells were seed into 96 well plate by 5000 per well, basal 1640 serum change preceding 10% FBS-1640 serum after 24 hours, Ramucirumab diluted by 5% FBS-1640 serum at a final concentration of 10^−3^-10^3^ μg/mL and change preceding culture serum. MTT assay detection after treatment for 24h. MKN45 cell is another poorly differentiated gastric carcinoma cells. 3-(4,5-dimethyl-2-thiazolyl)-2,5-diphenyl-2-H-tetrazolium bromide (MTT) assays evaluate each drugs’ inhibition ratio. Data statistics and significance analysis by OriginPro 8 and SPSS 19 sofware, four parameter logistic curve calculated by formula 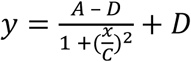, statistics C value (IC50) and R-square value.

### Cell fluorescence imaging

Ramucirumab was labeled by Alexa Fluor® 647 (Ex: 650 nm, Em: 670 nm, select Cy5 tunnel when imaging by conforcal and FCM) fluorescein firstly, utilaze Cy5 labelled antibody synthese Au-peg-Ab-Cy5. Au-peg-Ab-Cy5 or antibody-Cy5 treated SNU5 cells for 1, 2, 4 hours by final antibody concentration of 1 μg/mL, Hoechst 33342 (10 μg/mL, 200 μL, Ex: 405 nm, Em: 488 nm) was used as the nuclei indicator for 15 min incubation at 37 °C.

### In vivo fluorescence imaging

In order to detect the delivery efficiency of AuNR-peg-Ab in a real tumor system, *in vivo* imaging assay was implementation. Since Cy5 was Near infrared fluorescent dye, AuNR-peg-Ab-Cy5 could be utilized directly. All the *in vivo* experiments were in accordance with the *Beijing University Animal Study Committee*’s requirements and approved by the institutional ethical committee of the *National Center for Nanoscience and Technology*. we evaluated the delivery efficiency and the biodistribution of the nanoparticles for the BALB/c mice (Beijing Vital River Laboratory Animal Technology Co., Ltd) bearing SNU5 xenograft tumors by Cy5 imaging on Maestro *in vivo* spectrum imaging system. Collect 10^7^ SNU5 cells and inject into 50-56 day-age-BALB/c-nu mouse right hind leg for one week to forming a solid tumor. Each mouse received an intravenous injection of drugs which were pre-degermed by 0.22μm filter membrane. Mice were euthanized, the major organs (liver, lung, spleen, kidney, and heart) and tumor were dissected, and their fluorescence images were obtained.

### Flow cytometry analysis of cellular uptake Ab

The cellular uptake of FITC-labeled Ab and other nanodrugs were investigated by a flow cytometry analysis. cells were incubated with FITC-labeled peptides for 15min at 4 °C. Then these cells were washed with cold PBS and re-suspended in PBS for analysis. The FITC fluorescence intensity was measured with a flow cytometer (Becton Dickinson, USA).

### Protein digestion and LC-MS/MS analysis

This analysis method was done as we have previously described before^44^. Briefly, digestion of proteins from SNU5 cell was performed prior to LC–MS/MS analysis. Digested samples were analyzed by LC–MS/MS using an Easy-nLC1200 nanoflow UHPLC (Thermo Fisher Scientific Inc.) coupled to a Q Exactive-plus mass spectrometer. Data produced were searched using MaxQuant software (Computational Systems Biochemistry, Martinsried, Germany) package (version 1.5.1.2), against the SwissProt_2016_04 database with taxonomy of [human] selected.

### RNA isolation and sequencing

This analysis method was done as described before^45^. Briefly, total RNA was extracted in the cell. After RNA-sequencing, RNA-Seq reads were mapped to the human reference genome sequence (hg19, Genome Reference Consortium GRCh37) using Hisat2 (version 2.1.0), which uses HTSeq software and DEseq2 to determine the differentially expressed genes. And the transcript counts for gene expression levels were calculated, and the relative transcript abundance was determined as fragments per kilobase of exon per million fragments mapped (FPKM) using Cufflinks software (version 2.1.1).

### Bioinformatics analysis

Heat maps were produced using Perseus (1.6.2.2)^46^. Gene ontology and KEGG pathway analyses were performed using GeneCodis 3.0 ^47^. FDR (q value) was used to select interesting protein and gene sets.

### In vivo therapy assays of NR and SP group

About 6–8 weeks-old BALB/c mice were subcutaneously implanted SNU5 cells for therapy. the mice were randomly divided into three groups. When the tumors had been allowed to develop to approximately 100-200 mm^3^, mice were injected intravenously with AuNR-peg-Ab (NR group), AuSP-peg-Ab (SP group), Ab and PBS (100 µL) at a dose corresponding to 8 mg/kg of Ramucirumab (n=5). PBS was served as control. Administration was carried out on once a week considering the drug was teated in GC patient once two weeks. The tumor sizes and weights were recorded daily at the same time. Tumor sizes were measured by a vernier caliper. Tumor volume was calculated by the formula (L × W2)/2. L is for the longest and W is the shortest in tumor diameters (mm). The experimental data were assessed as the mean standard deviation using Origin software for four independent experiments. After that, tumors and other organs were collected. The toxicity in tumors and organs of each group were determined by H&E. The apoptosis of tumor cells was also detected by TUNEL assay.

### Real-time polymerase chain reaction (RT-PCR) and western blot assay for checking the expression of FcγR

Cells were cultured in the medium at the density of 10^7^ per dish. Total RNA was isolated using TRIzol reagent (Invitrogen) followed the manufacturer’s instructions. RNA was reversely transcribed using the first strand synthesis kit (TaKaRa) and cDNA was subjected to real-time PCR using SYBR green mix (TaKaRa). PCR cycles were performed on an ABI 7500 PCR System. Values obtained for the threshold cycle for each gene were normalized using the average of housekeeping genes GAPDH amplified on the same cycle. For western blot, cells were washed twice with ice-cold PBS and lysed in lysis buffer (Sigma, USA) on ice for 30 min. Cell lysates were collected with centrifugation at 4°C for 30 min and the total protein was measured using the BCA (bicinchoninic acid) protein determination assay kit (Beyotime Biotechnology, China).

According to the standard western blot procedures, proteins were separated by SDS-PAGE and transferred onto PVDF (polyvinylidene fluoride) membranes (Merck Millipore, DE). The membranes were then blocked for 1 h in 5% non-fat milk in PBST and incubated with CD64 and CD16 antibodies and β-actin antibody respectively overnight at 4°C. After washing 3 times with PBST for 5 min each, the membranes were incubated with horseradish peroxidase-coupled isotype-specific secondary antibodies for 1 h at room temperature. The immune complexes were detected by ChemiDoc MP imaging system (Bio-Rad, USA).

### Statistical analysis

All functional experiments were done in triplicate. Data were averaged and expressed as mean ± standard deviation (SD). They were analyzed with one-way analysis of variance (ANOVA) using SPSS software (version 20.0; IBM, New York, NY) and

## ASSOCIATED CONTENT

### Supporting Information

Detailed experimental materials and methods can be found in the Supporting Information.

