## supplemental materials for "Gold nanorod based delivery system could bring about superior therapy effect of Ramucirumab through direct cytotoxicity to cancer cell mediated by differential regulation of phagocytosis in gastric cancer cell"

**Supporting Information**

1 **Supplemental Materials**

**1.1 Ramucirumab lable Cy5 purification by HPLC.**

Ramucirumab was labeld by Cy5 using NHS wster labeling of an amino biomolecules kit. briefly, 100 μL 20 mg/mL antibody added 800 μL 0.1M NaHCO_3_ in 1.5 mL microcentrifuge tube, dissolve NHS ester (1 μg/mL) 66 μL into tube and vortex well, incubate the mixture overnight at room temperature. Purify the conjugate using gel-filtration method. SEC-HPLC analyze the product and check the sharp and retention time of prominent peak. **Figure S1** showed that retention time of Ab-Cy5 had been moved up (Molecular Weight was increased ) and shoulder peak appeared(Ab and Ab-Cy5 mixture).

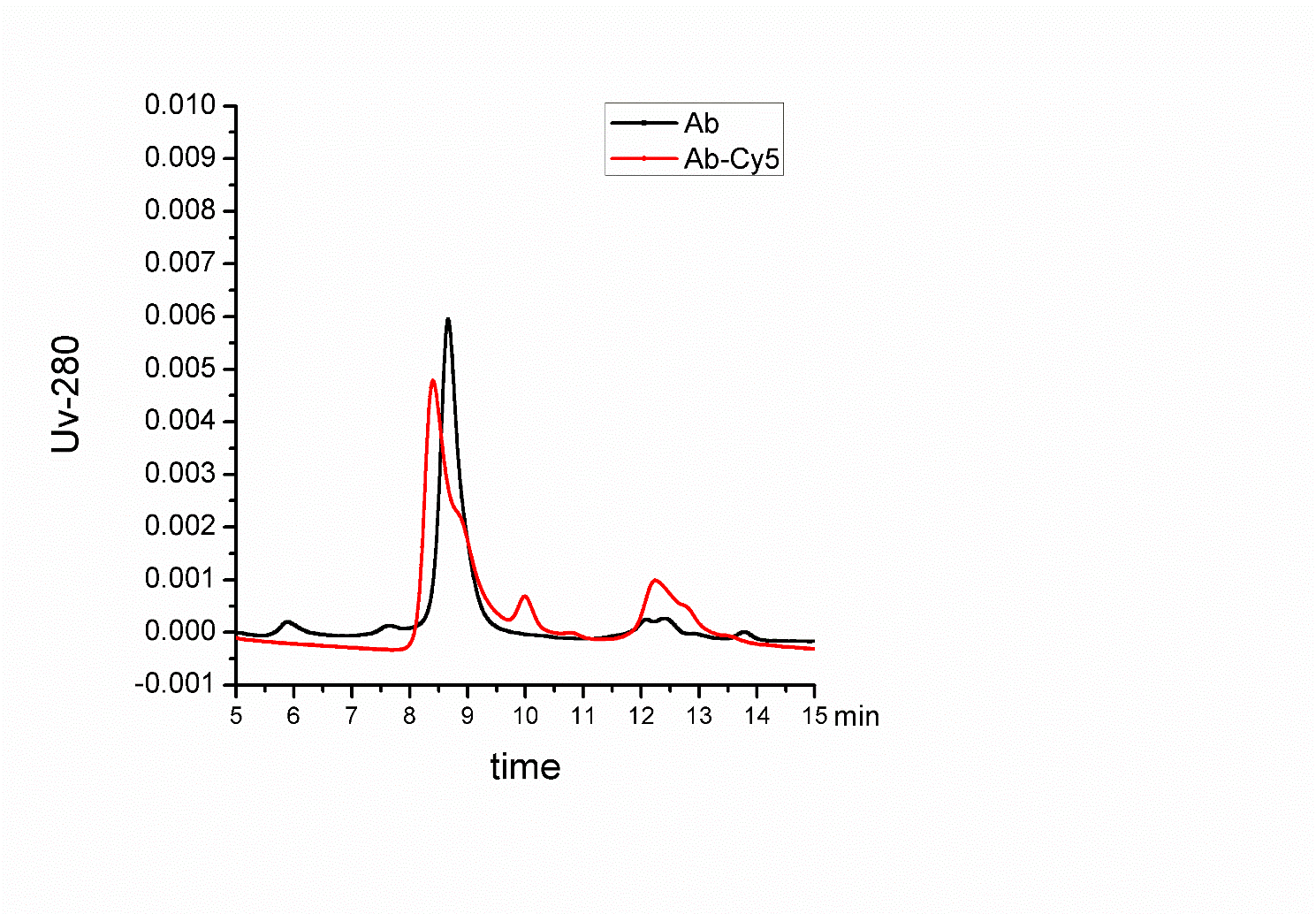

**Figure S1** SEC-HPLC analysis of Antibody and antibody labeled by Cy5

1.2 **Ellman DTNB method analyze.**

Standard calibration curve for PEG chains, whose concentration can be calculated via the following equation: Abs at 412nm = 0.0895×[PEG, mg/mL]+0.4967, R^2^=0.999. Variation of the excess of PEG thiolate chains as a function of the initial concentration in the incubation with 1 mL of nanorods. The dashed vertical line indicates the 100% saturation, i.e. the PEG concentration above which no more PEG can be bound to the nanoparticle’s surface.

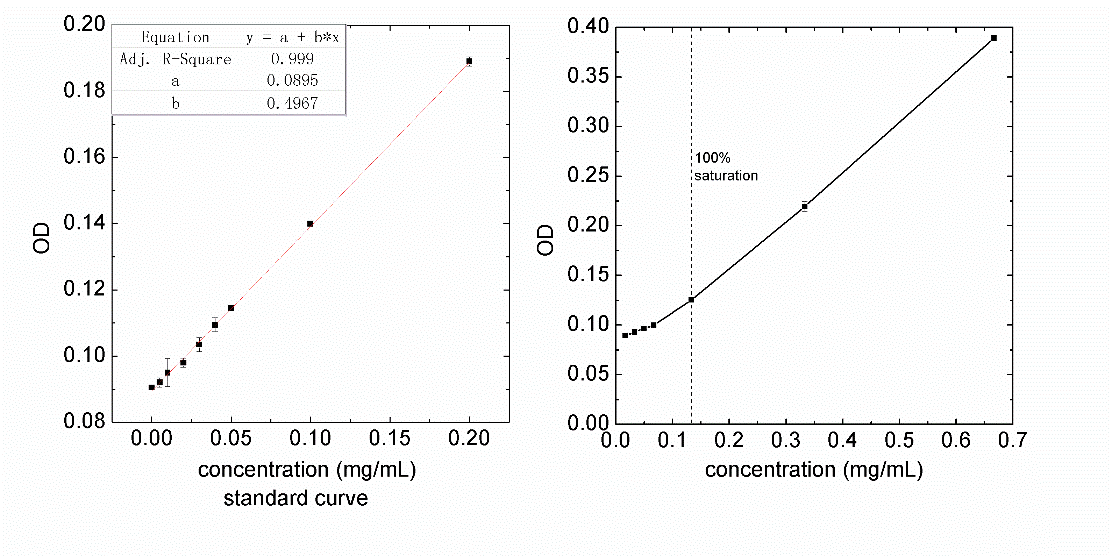

**Figure S2** Ellman’s Assay evaluate functionalization of gold nanorods with poly (ethylene glycol) (Peg)

**1.3 Selection for differential protein and expressed genes in the proteomics and transcriptomics.**

The analyses were based on diﬀerentially proteins and expressed genes in each treatment groups. Diﬀerential expression was defined as multiple testing adjusted p values smaller than or equal to 0.05 and fold change greater than or equal to 2.0-fold.

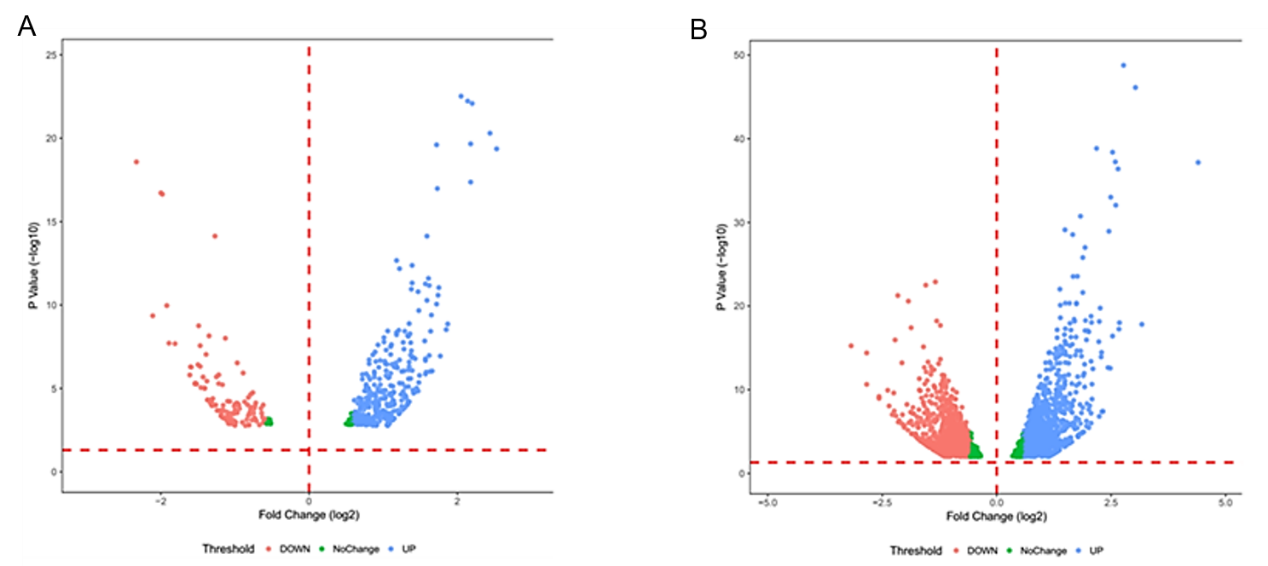

**Figure S3.** Volcano plots of Quantitative proteomics (A) and transcriptomics (B) results. The volcano plots were assembled which represented that the highly deregulated proteins and genes marked with red (SP group) or blue (NR group) color appear in the left or right sides. They showed the different expression genes for the different groups.

**1.4 Transcriptomics analysis of AuNR-peg-Ab (NR group) and AuSP-peg-Ab (SP group) compared with Ab treated group respectively in SNU5 cells.**

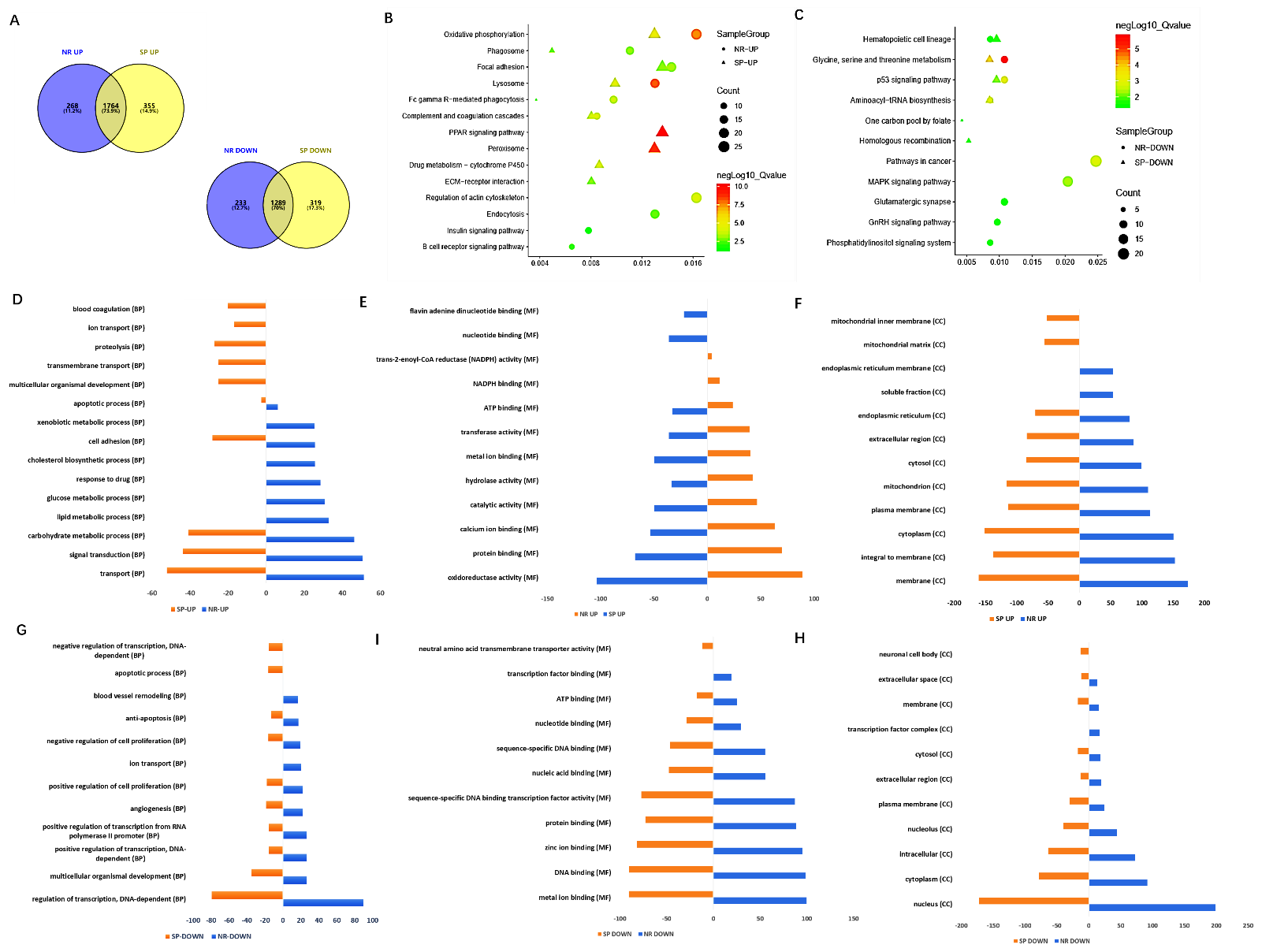

**Figure S4.** Transcriptomics analysis of AuNR-Peg-Ab (NR group) and AuSP-Peg-Ab (SP group) treated groups compared with Ab treated group respectively in SNU5 cell. A) Venn grams of the whole numbers of differential expressed genes quantified in transcriptomics results from different comparations. B-C) KEGG enrichment of differential genes. the color indicated the level of q value and the items with q<0.05 were included. D-F) Top 10 items in the Gene Ontology (GO) biological process (BP), molecular function (MF), cellular component (CC) enrichment of up-regulated genes in the different comparisons. G-H) Top 10 items in the Gene Ontology (GO) biological process (BP), molecular function (MF), cellular component (CC) enrichment of down-regulated genes in the different comparisons.

**1.5 Flow cytometry analysis of cellular uptake of Ab in different cell type.**

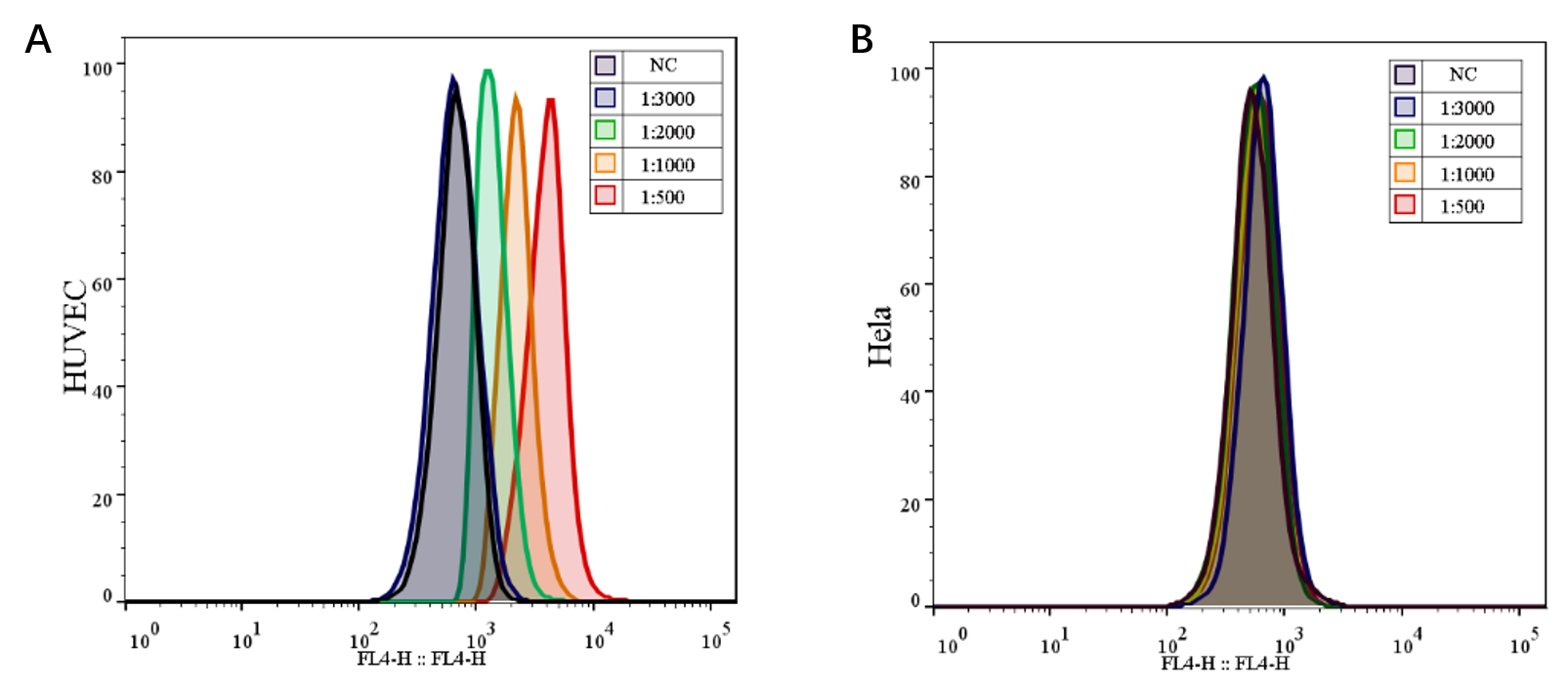

**Figure S5.** A-B), Flow cytometry analysis of Ab in HUVEC and Hela cells.

**2 Material and method**

**2.1 Design of gold particles-peg conjugate to Cy5-labeling of Ramucirumab and/or doxorubicin**

**2.8 Bioinformatics analysis**

Heat maps were produced using Perseus (1.6.2.2)[5]. Gene ontology and KEGG pathway analyses were performed using GeneCodis 3.0 [6]. FDR (q value) was used to select interesting protein and gene sets.

**2.11 Statistical analysis**

All functional experiments were done in triplicate. Data were averaged and expressed as mean ± standard deviation (SD). They were analyzed with one-way analysis of variance (ANOVA) using SPSS software (version 20.0; IBM, New York, NY) and the level of statistical significance was defined as p < 0.05
